# Age-Associated Decline in Autophagy Pathways in Retinal Pigment Epithelium and Protective Effects of Topical Trehalose in Light-induced Outer Retinal Degeneration in Mice

**DOI:** 10.1101/2025.01.14.632938

**Authors:** Katherine Cox, Gongyu Shi, Neve Read, Kepeng Ou, Zijia Liu, Jiahui Wu, Suci Cendanawati, Jenna Le Brun Powell, Lindsay B. Nicholson, Andrew D. Dick, Jian Liu

## Abstract

Age is a primary risk factor for chronic conditions, including age-related macular degeneration (AMD). Impairments in autophagy processes are implicated in AMD progression, but the extent of autophagy’s contribution and its therapeutic potential remain ambiguous. This study investigated age-associated transcriptomic changes in autophagy pathways in the retinal pigment epithelium (RPE) and evaluated the protective effects of topical trehalose, an autophagy-enhancing small molecule, against light-induced outer retinal degeneration in mice. Transcriptomic analysis of human RPE/choroid and mouse RPE revealed consistent downregulation of autophagy pathways with age, alongside variable changes as AMD severity progressed. Given the age- and AMD-associated perturbation of autophagy pathways, we examined trehalose treatment *in vitro*, which enhanced autophagic flux and restored mitochondrial respiratory function in primary murine RPE cells exposed to oxidative stress. *In vivo*, topical trehalose improved autophagy-lysosome activity in mouse RPE, demonstrated by elevated LC3B turnover and SQSTM1/p62 degradation. Furthermore, trehalose eyedrops protected mice from light-induced damage to the RPE and photoreceptors, preserving outer nuclear layer thickness, RPE morphology, and junctional F-actin organization. Taken together, the data support that age-related decline and severe dysregulation in autophagy contributed to AMD progression. By restoring autophagic flux, topical trehalose demonstrates therapeutic potential to address early autophagy-related pathological changes in AMD.

## 1. INTRODUCTION

Age-related macular degeneration (AMD) is a progressive multifactorial neurodegenerative disease that affects particularly the retina’s macular region, leading to severe central vision loss. While strong genetic associations, environmental factors and lifestyle choices all contribute to the complex disease pathogenesis, aging remains a significant risk factor (Fleckenstein, Schmitz-Valckenberg, & Chakravarthy, 2024). The retinal pigment epithelium (RPE), central for maintaining the visual cycle and photoreceptor health and outer retinal homeostasis, is particularly susceptible to aging not least because of the constant exposure to high levels of oxidative stress. The RPE shows the highest number of differentially expressed genes (DEGs) in the retina associated with aging, and its degeneration is a hallmark of AMD (S. Wang et al., 2020). Early AMD results in the deposition of oxidized lipoproteinaceous drusen beneath the RPE, and may progress to late forms, including advanced dry AMD (geographic atrophy, GA) or, in 10-15% of cases, wet AMD (neovascular). Effective treatments to halt the degenerative process of AMD are awaited, despite current anti-complement therapies for GA that offer limited functional benefits (Csaky, Miller, Martin, & Johnson, 2024).

Proteomic analysis reveals that the protein profile of drusen closely resembles that of the RPE, indicating that degenerating RPE is a primary source of drusen components, likely through exosome-mediated release (Crabb, 2014; A. L. Wang et al., 2009). This suggests a disruption in the RPE’s clearance pathways, such as autophagy, a catabolic process required by RPE because of its high metabolic rate. During autophagic flux, dysfunctional organelles and cytotoxic macromolecular aggregates are degraded through the formation of autophagosomes and the consequent fusion with lysosomes (autolysosomes) (Menzies et al., 2017). Under conditions of nutrient deprivation or energy stress, autophagic cascade is triggered by the inhibition of the mammalian target of rapamycin (mTOR) and activation of 5’-adenosine monophosphate-activated protein kinase (AMPK), respectively (Menzies et al., 2017).

Impaired autophagy has been linked to various age-related conditions, including neurodegeneration, type 2 diabetes, and cancer (Menzies et al., 2017). Conversely, enhanced autophagy may serve as a defence mechanism against cellular damage in certain disease contexts by increasing the turnover of abnormal or excessive cellular components (Y. Wang et al., 2013). In AMD, transcriptomic analyses of RPE cells from the macular region reveal a significant upregulation of genes involved in autophagy initiation (Orozco et al., 2023; Ramírez-Pardo et al., 2023), suggesting an adaptive response of the remaining RPE cells to counteract the stressors driving RPE dysfunction. Paradoxically, despite this transcriptomic upregulation, AMD progression is characterized by a functionally reduced autophagy flux (Kaarniranta et al., 2023; Vessey et al., 2022). Dysfunctional autophagy is associated with increased transcytosis and exocytosis, heightened susceptibility to mitochondrial oxidative stress, elevated mitochondrial damage, disrupted proteostasis, inflammasome activation, and drusen formation (J. Liu et al., 2016; Ramírez-Pardo et al., 2023).

Autophagy is considered as a promising therapeutic target due to its amenability to chemical modulation. Small molecules (MW < 900 Daltons) with compact size (∼1 nm) can efficiently penetrate cells and access intracellular targets via passive or carrier-mediated transport (Ramsay et al., 2023). One such molecules is trehalose (MW = 342.3 Daltons) which is a naturally abundant, nonreducing disaccharide of D-glucose. Although trehalose is not endogenously produced in mammals and is hydrophilic, it penetrates mammalian cell membranes via fluid-phase endocytosis at millimolar concentrations (Huang et al., 2024; Zhang, Oldenhof, Sieme, & Wolkers, 2017). Trehalose has shown autophagy-dependent therapeutic effects against mitochondrial dysfunction, inflammation, cardiac disease, and neurodegeneration (Korolenko et al., 2021; Sciarretta et al., 2018). Mechanistically, trehalose enhances autophagy through multiple pathways, including activating AMPK signaling, incorporating into autophagosomes, and stimulating lysosomal activity (Rusmini et al., 2019; Xu, Chen, Sheng, & Yang, 2019). Widely used in 3% w/v (87.7 mM) ophthalmic formulations, trehalose effectively treats dry eye disease by reducing conjunctival inflammation, restoring osmotic balance, and protecting epithelial cells (Cagini et al., 2021). Its low-toxicity profile supports topical administration at concentrations as high as 30% (Laihia & Kaarniranta, 2020), highlighting its potential for high-dose or long-term applications. Alongside ongoing clinical trials, preclinical studies on small molecule-based topical therapies have demonstrated significant advantages as non-invasive treatment options for retinal pathologies (Batson et al., 2017; Löscher, Seiz, Hurst, & Schnichels, 2022).

Here, we explored age-related transcriptomic changes in human and murine RPE and/or choroidal samples, focusing on autophagy and related pathways. We conducted an interspecies comparison to give provenance to examine the effects of trehalose on autophagy flux, mitochondrial oxidative stress, and respiratory capacity in primary murine RPE cells. Additionally, we evaluated the therapeutic efficacy of topical trehalose in protecting against RPE and photoreceptor damage in light-induced outer retinal degeneration in mice.

## 2. RESULTS

### 2.1 Autophagy, mitophagy and lysosome pathways are predominantly downregulated in human RPE/choroid with age

To identify potential mechanisms contributing to the onset of AMD, we extracted transcriptomic data of non-AMD individuals from a recent study to analyze age-associated changes in human RPE/choroid samples (Orozco et al., 2023). Due to the lack of young donor eyes in the dataset, Orozco *et al*. conducted a linear analysis of macular RPE/choroid samples from 36 normal control donors aged 50 to 94 years (private communication), with a mean age of 70.5 ± 13.6 years and a mixed-gender cohort (Orozco et al., 2023). The original study identified 789 downregulated and 725 upregulated genes correlated with increasing age (Data S1, and Figure S2a,b), without further stratification. Leveraging this dataset, we performed pathway analyses to explore transcriptomic alterations associated with aging.

As shown in Figure 1a, the top Kyoto Encyclopedia of Genes and Genomes (KEGG) pathways associated with age-related downregulated DEGs were linked to cellular transport and signaling (Endocytosis, Rap1 signaling, MAPK signaling, Apelin signaling, and Axon guidance), cellular degradation and recycling (Proteolysis, Lysosome, and Mitophagy), metabolic pathways (Fatty acid metabolism, N-glycan biosynthesis, and Sphingolipid signaling), and cellular stress and protein synthesis (Chemical carcinogenesis - reactive oxygen species (ROS), and Ribosome). In contrast (Figure 1b), upregulated genes were significantly enriched in structure and adhesion pathways (Cytoskeleton, and Tight junction), protein and lipid metabolism (Protein digestion and absorption, Lysine degradation, and Glycerophospholipid metabolism), signaling pathways (Notch, Hedgehog, and mTOR signaling), and disease-associated pathways (Dilated cardiomyopathy, and Alcoholic liver disease). These results indicate a mixed regulation of cellular signaling and metabolic pathways, and patterns of downregulation in Mitophagy and Lysosome pathways.

**Figure 1:**
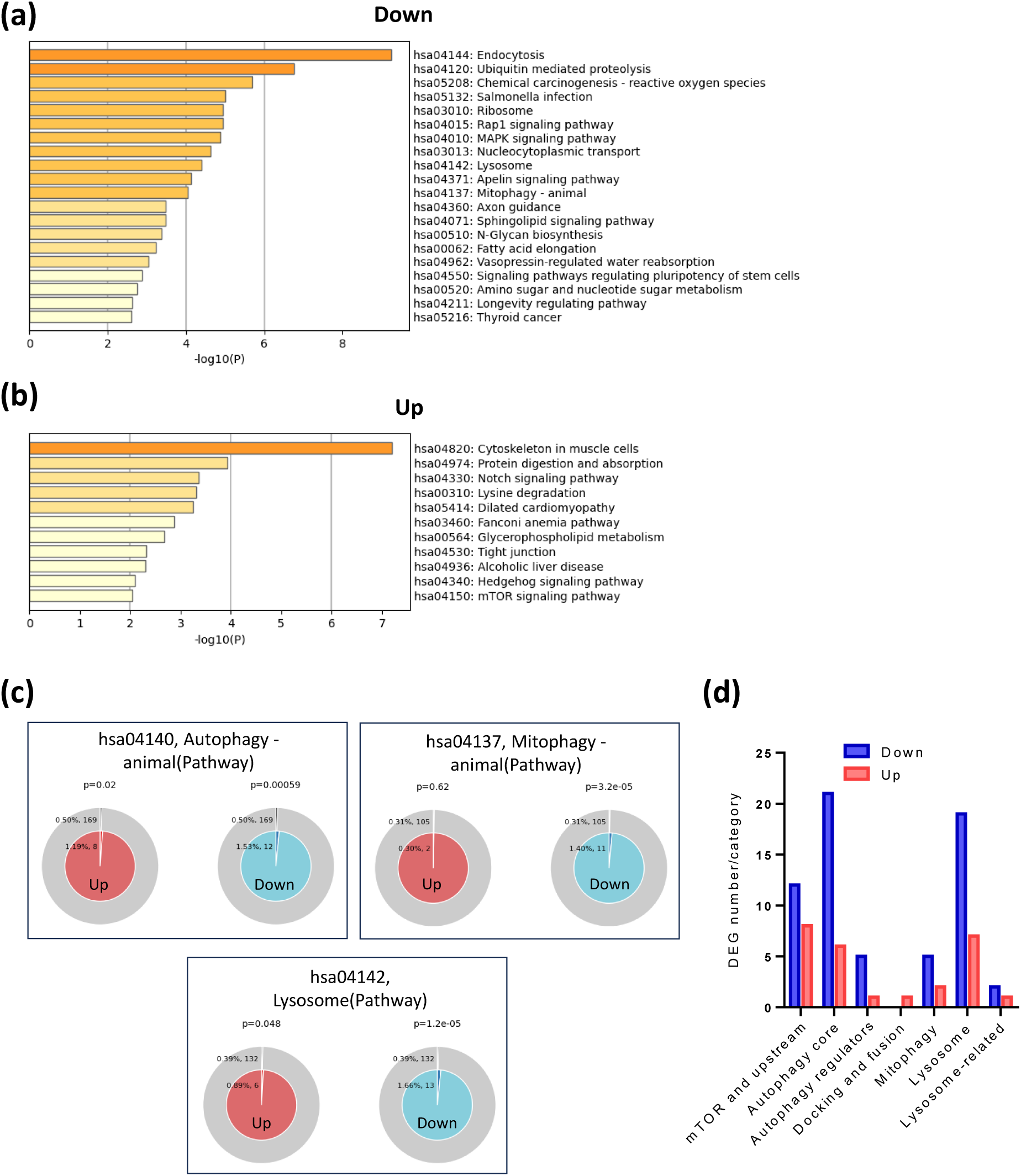
Predominant downregulation of autophagy-lysosomal pathways in aged human RPE/choroid. (**a** and **b**) Top enriched KEGG pathways of downregulated (**a**) and upregulated (**b**) genes identified through a linear analysis of RNA-seq data from macular RPE/choroid samples of 36 normal controls aged 50-94 years (mean 70.5 ± 13.6, mixed gender) (Orozco et al., 2023). Pathway enrichment analysis was conducted using the Metascape platform with default setting: P cutoff = 0.01, minimum overlap = 3, and minimum enrichment = 1.5. (**c**) Membership analysis conducted through Metascape for upregulated (left) and downregulated (right) genes enriched in specific KEGG pathway terms, including Autophagy (hsa04140), Mitophagy (hsa04137), and lysosome (hsa04142). The outer circle represents the percentage and number of total genes associated with the pathway, while the inner circle shows the percentage and number of input genes mapping to the pathway. The *P*-value above the pie chart indicates the statistical significance of the gene set membership to the pathway term. (**d**) Number of downregulated (blue) and upregulated (red) genes in each category based on a broader autophagy-lysosome gene list (Bordi et al., 2021).

Membership analysis further revealed that the Autophagy pathway displayed predominant downregulation with age (P = 0.00059), alongside a significant albeit minor upregulation (P = 0.02) (Figure 1c). Similarly, pronounced age-related downregulation was confirmed in Mitophagy (P = 3.2e-5) and Lysosome pathway (P = 1.2e-5), while a minor upregulation in Lysosome pathway was also noted (P = 0.048).

To further elucidate the extent of transcriptomic dysregulation in autophagy-lysosome pathways with age, we analyzed the lists of downregulated and upregulated genes in conjunction with a previously identified autophagy-lysosome gene set (Bordi et al., 2021). This gene set covers a broader range of 604 genes and more diverse categories based on their roles within these pathways, compared to the KEGG terms Autophagy (hsa04140) and Lysosome (hsa04142), which only includes 169 and 132 genes, respectively. Consistent with the pathway analysis, we found a greater number of downregulated genes across most autophagy pathway categories, except for Docking and fusion (Figure 1d). Both downregulated and upregulated genes predominantly aligned with mTOR and upstream components, Autophagy core machinery, and Lysosome pathways, suggesting significant dysregulation in these critical categories (Figure 1d). A full list of DEGs overlapped with the broader autophagy-lysosome pathway categories is provided in Data S2.

### 2.2 Autophagy and mitophagy pathways are predominantly downregulated in mouse RPE with age, resembling human data

We next evaluated whether transcriptomic changes in autophagy and related pathways in aging mouse eye tissue reflect those observed in humans, focusing specifically on the RPE due to its high autophagic demands and susceptibility to aging (Intartaglia, Giamundo, & Conte, 2022). Male mice were used to minimize sex-dependent variations in autophagy (Bjornson, Vanderplow, Bhasker, & Cahill, 2024). RNA-seq analysis was performed on murine RPE cells isolated from 5- and 22-month-old mice, equivalent to human ages of 20-30 and 70-80 years, respectively (Graff, Payne, & El-Bouri, 2021). The experiments were conducted in two batches with totally 6 eyes per group. The sample information is provided in Data S3. After batch correction, normalization, and exclusion of low read counts (basemean ≤ 20), the gene dataset (Data S3) underwent quality control using iDEP.96, confirming comparable distribution of transformed expression values across samples (Figure S1a). Minimal variation within each age group and clear separation between the two age groups were confirmed by Principal Component Analysis (PCA) (Figure S1b). Raw and processed gene counts are listed in Data S3.

DESeq2 analysis identified 1412 downregulated and 1694 upregulated DEGs (Padj < 0.05, |log2FC| > 0.25) in aged RPE compared to young controls (Figure 2a and Data S3). Enrichment analysis using the PaGenBase dataset via Metascape showed that both downregulated and upregulated DEGs were predominantly associated with the RPE (Figure 2b). The upregulated genes were also strongly associated with tissues specific to eye, retina, or brain. This finding validated the purity of the RPE samples, supporting the Western blot analysis showing RPE65-positive and rhodopsin-negative fractions as we reported elsewhere (Scott et al., 2021).

**Figure 2:**
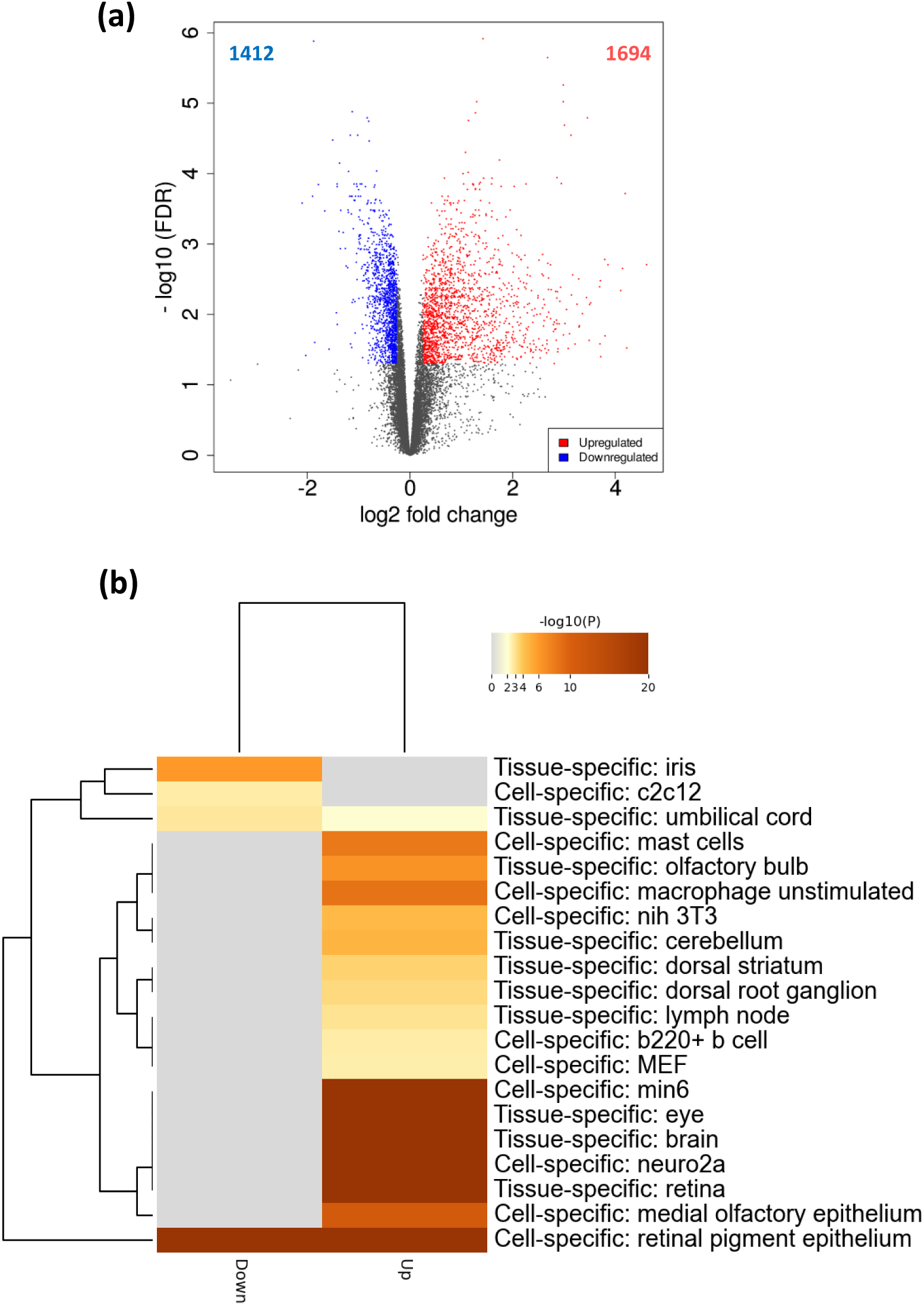
Volcano plot and PaGenBase enrichment of DEGs in aged mouse RPE. RPE cells isolated from eyes of 5- and 22-month-old male C57BL/6J mice (n = 6 eyes per group) were subjected to RNA-seq analysis. (**a**) The volcano plot displays significantly downregulated (blue) and upregulated (red) DEGs, with the number of DEGs indicated. Statistical thresholds were set at Padj < 0.05 and |log2FC| > 0.25. Genes without significant changes are presented in grey. (**b**) PaGenBase enrichment analysis, performed via Metascape, illustrates cell and tissue specificity of downregulated and upregulated DEGs. The entire genome was used as the enrichment background. Terms with P cutoff = 0.01, minimum overlap = 3, and minimum enrichment = 1.5 were identified and grouped into clusters based on membership similarities.

As shown in Figure 3a, the top KEGG-enriched terms for downregulated DEGs in aged mouse RPE include pathways related to cellular signaling and regulation (Rap1 signaling, Apelin signaling, PPAR signaling, Focal adhesion, and Gap junction), metabolism and biosynthesis (Fatty acid metabolism, Regulation of lipolysis in adipocytes, and Tyrosine metabolism), cellular processes and degradation (Mitophagy, Peroxisome, Protein processing in endoplasmic reticulum, Axon guidance, and Salmonella infection), and disease-related pathways (Chemical carcinogenesis - ROS, Pathway in cancer, and Fluid shear stress and atherosclerosis). Upregulated DEGs are enriched in pathways involving nervous system and synaptic function (Synaptic vesicle cycle, GABAergic synapse, Phototransduction, Nicotine addiction, and Axon guidance), cellular signaling and communication (Calcium signaling, Chemokine signaling, MAPK signaling, Cell adhesion molecules, ECM-receptor interaction), immune responses (Natural killer cell-mediated cytotoxicity, Primary immunodeficiency, and Fc gamma R-mediated phagocytosis), and metabolic and immune-related disease pathways (Dilated cardiomyopathy, Type II diabetes mellitus, and Rheumatoid arthritis).

**Figure 3:**
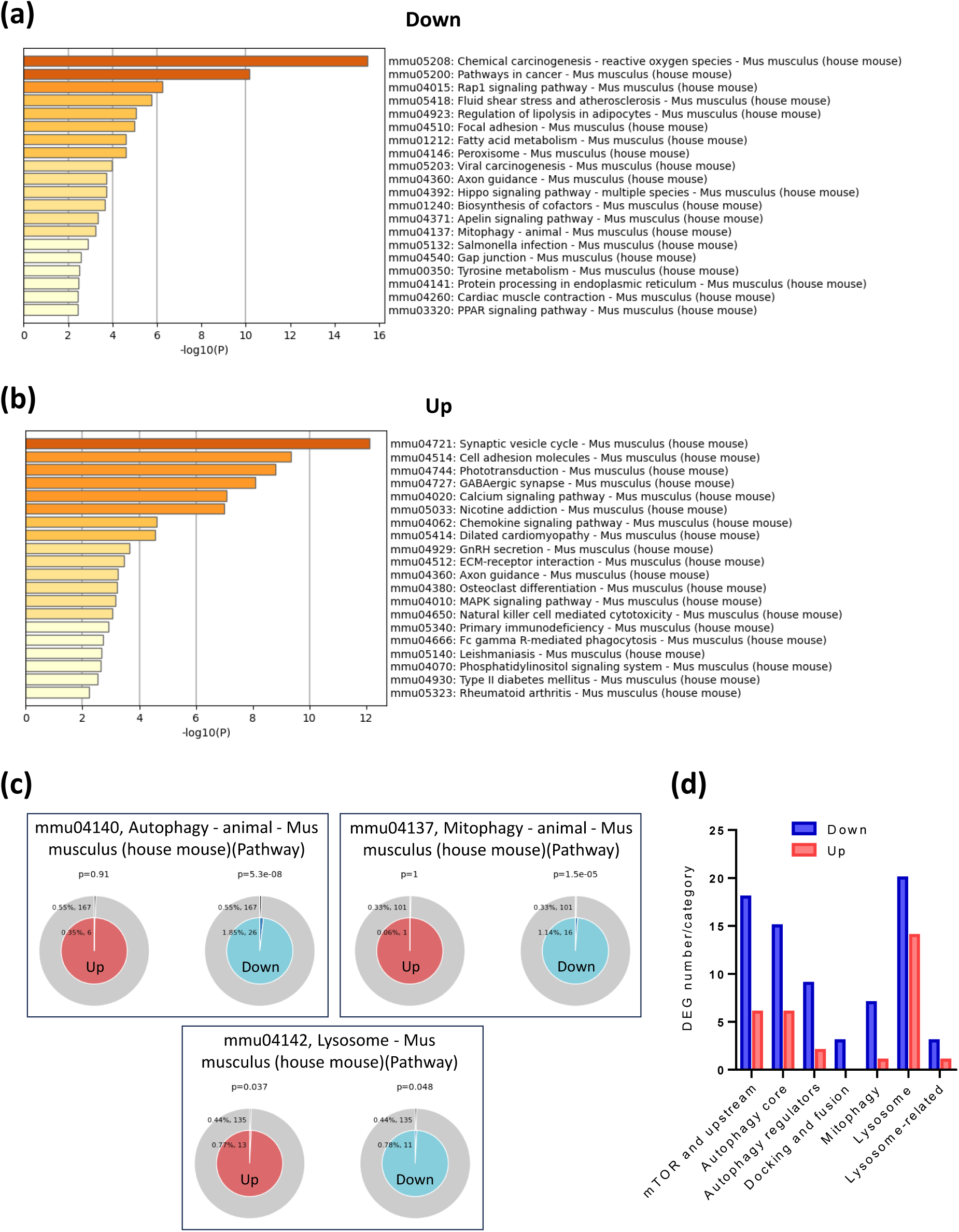
Predominant downregulation of autophagy-lysosomal pathways in aged mouse RPE. (**a** and **b**) Top enriched KEGG pathways of downregulated (**a**) and upregulated (**b**) DEGs identified through RNA-seq analysis of RPE cells isolated from 5- and 22-month-old male C57BL/6J mice (n = 6 per group). Pathway enrichment analysis was conducted using the Metascape platform with default setting: P cutoff = 0.01, minimum overlap = 3, and minimum enrichment = 1.5. (**c**) Membership analysis conducted through Metascape for upregulated (left) and downregulated (right) genes enriched in specific KEGG pathway terms, including Autophagy (mmu04140), Mitophagy (mmu04137), and lysosome (mmu04142). The outer circle represents the percentage and number of total genes associated with the pathway, while the inner circle shows the percentage and number of input genes mapping to the pathway. The P-value above the pie chart indicates the statistical significance of the gene set membership to the pathway term. (**d**) Number of downregulated (blue) and upregulated (red) genes in each category based on a broader autophagy-lysosome gene list (Bordi et al., 2021).

Between mouse RPE and human RPE/choroid, the Mitophagy pathway was one of the consistently downregulated pathways with age in both species. Other overlapping downregulated pathways include Chemical carcinogenesis - ROS, Rap1 signaling, Fatty acid metabolism, Apelin signaling, and Salmonella infection (Figure 1a and Figure 3a), while overlapping upregulated pathways include Dilated cardiomyopathy (Figure 1b and Figure 3b). However, certain pathways showed opposite trends between species. For example, MAPK signaling was downregulated in aged human RPE/choroid (Figure 1a) but upregulated in aged mouse RPE (Figure 3b). Additionally, the Axon guidance pathway showed downregulation in aged human samples (Figure 1a), whereas it was both upregulated and downregulated in mice (Figure 3a,b).

Comparable to the human RPE/choroid data (Figure 1c), the Membership analysis of mouse RPE showed a significant downregulation in Autophagy (P = 5.3e-8) and Mitophagy (P = 1.5e-5) pathways (Figure 3c). The Lysosome pathway in mice exhibited mixed regulation (upregulation: P = 0.037; downregulation: P = 0.048).

The connection between dysregulated genes and autophagy pathways was next cross-referenced with the broader autophagy-lysosome gene list (Bordi et al., 2021). Similarly to human data (Figure 1d), DEGs associated with autophagy-lysosome were overwhelmingly downregulated across all categories (Figure 3d), and the majority of both downregulated and upregulated DEGs were involved in mTOR signaling, autophagy core, and lysosome pathways (Figure 3d and Data S4).

### 2.3 Transcriptomic changes in aging human RPE/choroid and mouse RPE exhibit significant overlap, particularly in downregulated pathways

Using Metascape (Zhou et al., 2019), we conducted a multiple gene list analysis to compare the correlation and diversity of transcriptomic signatures of aging between humans and mice. As shown in Figure S2a and b, 108 genes were downregulated in both human RPE/choroid (out of 789 total downregulated genes) and mouse RPE (out of 1438) with age. Meanwhile, 65 genes were upregulated in both humans (out of 725) and mice (out of 1760). A complete list of overlapping genes is provided in Data S5.

KEGG pathway enrichment analysis revealed that the overlapping downregulated genes between the two species were significantly associated with pathways including Fatty acid metabolism and degradation, Mitophagy and Autophagy, Endocytosis, Axon guidance, Hippo signaling, Cellular responses to Salmonella, and various cancer-related pathways (Figure S2c). In contrast, fewer overlapping pathways were identified in the upregulated genes, which were primarily related to Cytoskeleton, Protein digestion, and Dilated cardiomyopathy (Figure S2d).

Membership analysis unveiled that several key transcriptomic pathways associated with aging were shared between humans and mice (Figure S3). Pathways linked to RPE melanogenesis (Figure S3a,b), cellular stress response (Figure S3c,d), and mitochondrial respiratory function (Figure S3e,f) were similarly downregulated in both species, whereas bidirectional regulations were observed in immune system pathways (Figure S3g,h). The findings that age-associated regulatory patterns of critical pathways are common to human and mouse samples reinforce the value of mouse models for studying aging-related changes in the RPE.

### 2.4 Autophagy and lysosome pathways show stage-specific transcriptomic changes in human RPE/choroid during AMD progression

We next conducted Membership analyses for Autophagy (hsa04140) and Lysosome (hsa04142) pathways to investigate stage-specific alterations in autophagic processes associated with early AMD and throughout the progression of the disease. These analyses utilized the same transcriptomic dataset as above (Orozco et al., 2023), which is uniquely suited to our study for stage-specific analysis due to the relatively robust sample size, providing sufficient representation of each AMD severity stage. Comparisons were made for each stage of dry AMD (dAMD), including early (eAMD, n = 16), intermediate (iAMD, n = 8), as well as geographic atrophy (GA, n = 10), and neovascular AMD (nAMD, n = 18), against normal controls (n = 36). Additionally, a linear analysis was conducted to assess transcriptomic changes associated with dAMD progression from eAMD to GA. The stage-specific and progression-correlated DEGs are listed in Data S1.

Compared to normal controls, the Autophagy pathway showed no significant changes in eAMD. However, it was notably upregulated in iAMD (P = 9.1e-5), suggesting an adaptive cellular response to mitigate increasing stress. In advanced stages, mixed regulation was observed, with evidence of both upregulation (GA: P = 0.0018; nAMD: P = 7.2e-5) and downregulation (GA: P = 0.0051; nAMD: P = 0.011), indicating a significant perturbation of autophagic activity (Figure S4a). This is not surprising, as these late AMD stages are characterized by extensive cell death in the RPE and excessive cellular stress imposed within the remaining, compromised cells. Similarly, a linear analysis of dAMD progression from eAMD to GA revealed concurrent upregulation (P = 1.9e-6) and downregulation (P = 0.0031) of the Autophagy pathway (Figure S4b).

Stage-specific analysis of the Lysosome pathway unveiled mixed regulation in eAMD (upregulation: P = 0.0019; downregulation: P = 0.03) and nAMD (upregulation: P = 0.0024; downregulation: P = 0.027). In contrast, iAMD and GA exhibited predominant upregulation of Lysosomal pathway (P = 5.5e-18 and P = 0.00019, respectively) (Figure S4c). The Lysosome pathway showed predominant upregulation during progression of dAMD (P = 3.1e-8, Figure S4d).

To corroborate these changes in Autophagy and Lysosome pathways, we analyzed the Immune system pathway (R-HAS-168256), which demonstrated consistent and significant dysregulation across all stages of dAMD and nAMD, with a predominant trend toward upregulation (Figure S4e,f). Similarly, the MAPK signaling pathway (hsa04010) exhibited an overall upregulation across stages of AMD (Figure S4g,h).

These findings highlight dynamic, stage-specific transcriptomic alterations that occur during the onset and progression of AMD, emphasizing the complexity of molecular changes underlying disease development.

### 2.5 Trehalose stimulates autophagy flux and mitigates hydrogen peroxide-induced oxidative stress and mitochondrial dysfunction in mouse RPE cells

The analysis thus far demonstrated that genes related to autophagy pathways were dominated by a downregulation in the aged RPE of both humans and mice, exhibited minimal alterations in eAMD, and showed mixed but more elevated levels in the later stages of AMD. These results infer age-associated disturbances in autophagy that are likely associated with AMD development. We reverted to murine models to assess whether enhancing autophagic activity could prevent or treat oxidative stress-induced outer retinal degeneration. To start, we examined the effects of autophagy enhancement via trehalose *in vitro*, based on reported effective doses of trehalose (1 to 100 mM) in various cell types, including human primary or iPSC-derived RPE cells (Fisher et al., 2022). The toxicity of the treatment in primary murine RPE cells was assessed through a Lactate Dehydrogenase (LDH) assay and Mito Stress test. Our results demonstrated that trehalose concentrations up to 12.5 mM did not compromise cell membrane integrity or mitochondrial oxygen consumption rate (OCR) in the tested cells (Figure S5a-c), and trehalose at 12.5 mM was therefore adopted for modulating autophagy in primary RPE cells.

To assess autophagy flux, we used tandem-tagged dual fluorophore probes, GFP (green) and RFP (red), to track the maturation of autophagosomes and the formation of autolysosomes following the fusion to lysosomes. In untreated cells (Figure 4a, top), the RFP-GFP-microtubule-associated protein 1 light chain 3 beta (LC3B) tandem tracker unveiled autophagosomes as colocalised RFP and GFP (GFP+RFP+; yellow). As autophagosomes fuse with lysosomes, the acid-sensitive GFP diminishes, distinguishing autolysosomes (GFP-RFP+; red). Further validation was achieved by colocalizing RFP with Lysotracker (magenta). In cells with inhibited autophagy, such as those treated with chloroquine (CQ) (Figure 4a, middle), which blocks autophagosome-lysosome fusion and slows lysosomal acidification, autophagy flux was obstructed, resulting in autophagosome accumulation and a much more prominent GFP signal compared to RFP.

**Figure 4:**
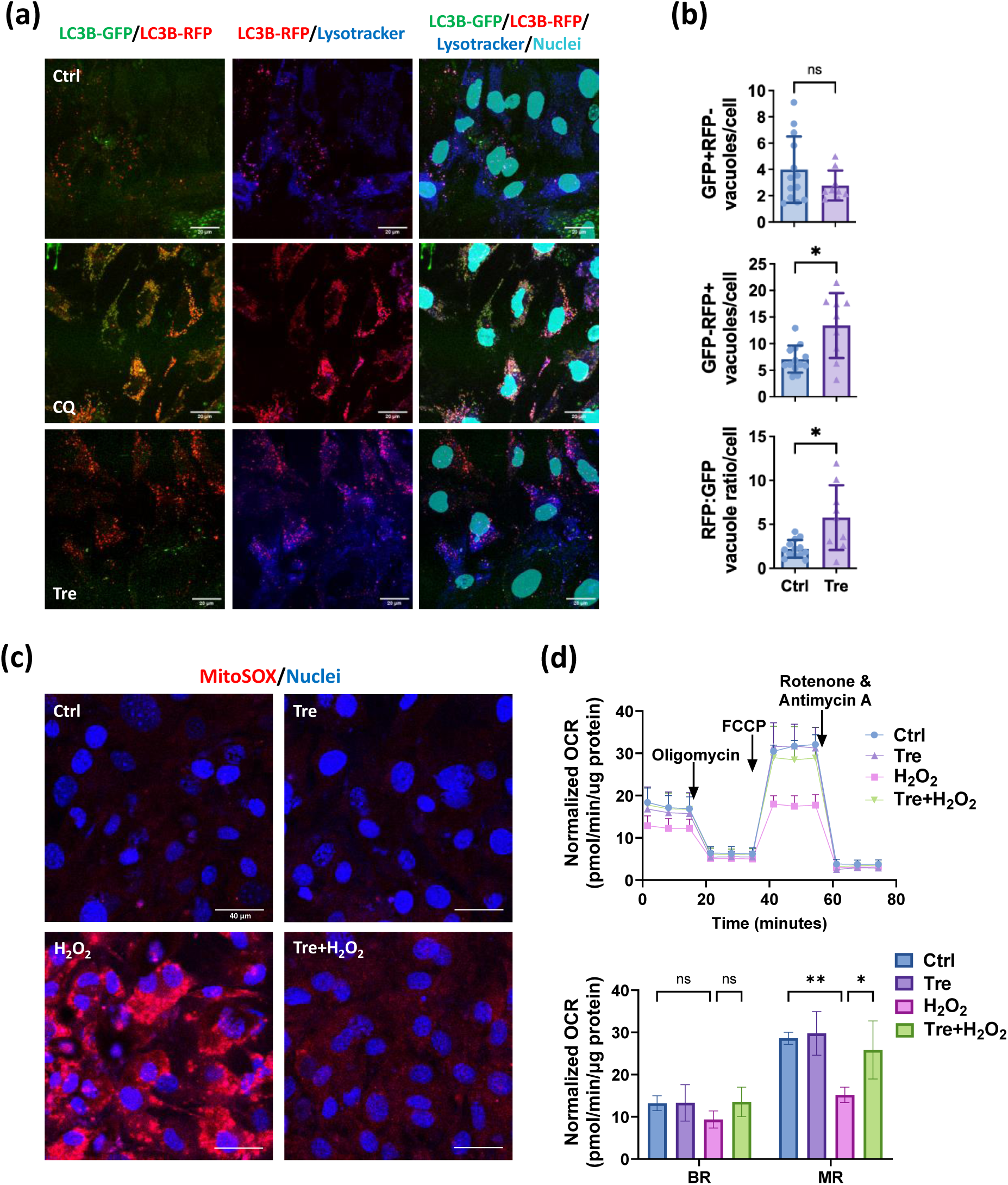
Trehalose effects in inducing autophagy flux and protecting mitochondrial activity against oxidative stress in primary murine RPE cells. (**a**) Confocal images of untreated RPE cells (top), and RPE cells treated with chloroquine (CQ, 50 µM, middle) or trehalose (12.5 mM, bottom) showing LC3B tandem tracker-stained autophagosome (LC3B-GFP, Green) and autolysosome (LC3B-RFP, Red), Lysotracker-stained lysosome (Blue) and Hoescht-stained nuclei (Cyan). (**b**) Quantification of autophagosomes (GFP+RFP-vacuoles), autolysosomes (GFP-RFP+ vacuoles), and autophagy flux (RFP:GFP ratio, bottom) in trehalose-treated RPE cells (n = 9), compared to control cells (n = 14). (**c**) Confocal images of MitoSox staining (red) in untreated RPE cells (top left), and RPE cells treated with trehalose (12.5 mM, top right), H_2_O_2_ (60 µM, bottom left), or a combination of both (bottom right). Hoescht staining in blue. (**d**) Oxygen consumption rate (OCR) profile (top) and basal respiration (BR) and maximal respiration (MR) parameters (bottom) of RPE cells treated with trehalose, H_2_O_2_, or their combination, compared to untreated cells (n = 3). Comparison by Mann Whitney test for GFP+RFP-(nonnormal data), Welch’s tests for GFP-RFP+ and RFP:GFP (normal data with unequal variances) (**b**), or two-way ANOVA with Bonferroni tests for OCR parameter analysis (**d**). *P < 0.05; **P < 0.01; ns = nonsignificant.

In contrast, treatment with trehalose in RPE cells led to an increase in the RFP signal colocalized with Lysotracker, indicating augmented autophagy flux (Figure 4a, bottom). As shown in Figure 4b, no difference was observed in the number of GFP+RFP-vacuoles, indicating efficient autophagosome-lysosome fusion. Trehalose significantly increased the number of autolysosome (GFP-RFP+) vacuoles, as well as the RFP:GFP ratio (Figure 4b), confirming a substantial enhancement of autophagy flux.

Using an *in vitro* model of acute oxidative stress induced by hydrogen peroxide (H_2_O_2_), we found that enhancing autophagy flux in RPE cells with trehalose effectively attenuated H_2_O_2_-induced cytoplasmic production of mitochondrial superoxide (MitoSOX, Figure 4c). Moreover, the OCR of the stressed cells, with or without the presence of trehalose, was measured before and after the addition of an ATP synthase inhibitor oligomycin and an electron transport chain uncoupler FCCP. While basal respiration (BR) of the mitochondria remained unchanged, H_2_O_2_ markedly reduced the maximal respiration (MR), which was significantly restored by trehalose (Figure 4d).

### 2.6 Topical trehalose application enhances RPE autophagic activity in mice

As trehalose enhanced autophagy flux and protected against oxidative stress in mouse primary RPE cells, we evaluated its effect *in vivo* via topical application. In our study, twice-daily administration of trehalose eyedrops at 3% (87.7 mM) or 9% (263.1 mM) concentrations (10 µl in PBS) over 2 weeks showed no significant side effects on the ocular surface or retina in treated mice (Figure S6). To maximize trehalose accessibility to the posterior segment of the eye, the higher dose was selected for subsequent experiments.

Autophagy flux after 2 weeks of treatment was determined by immunofluorescence co-staining of retinal sections with the autophagosome marker LC3B and the lysosome marker lysosomal-associated membrane protein 1 (LAMP1). Compared to the PBS-treated control, increased LC3B immunopositivity (red) was detected in the inner segment (IS) and RPE layers of the trehalose-treated eyes, along with a pronounced increase in LAMP1 signal (blue) in the RPE (Figure 5a). Colocalization of LC3B and LAMP1 signals (purple) in the RPE further confirmed enhanced autolysosome formation. Additionally, sequestosome-1 (SQSTM1)/p62 (from hereon p62), an intracellular waste cargo protein and marker of autophagic degradation that decreases with increased autophagy activity, was reduced in trehalose-treated eyes compared to PBS controls (Figure 5b). Quantitative analysis of mean fluorescence intensity (MFI) demonstrated significant increases in LC3B and LAMP1 (Figure 5c), and a decrease in p62 immunopositivity (Figure 5c) in the RPE of trehalose-treated eyes compared to PBS-treated controls.

**Figure 5:**
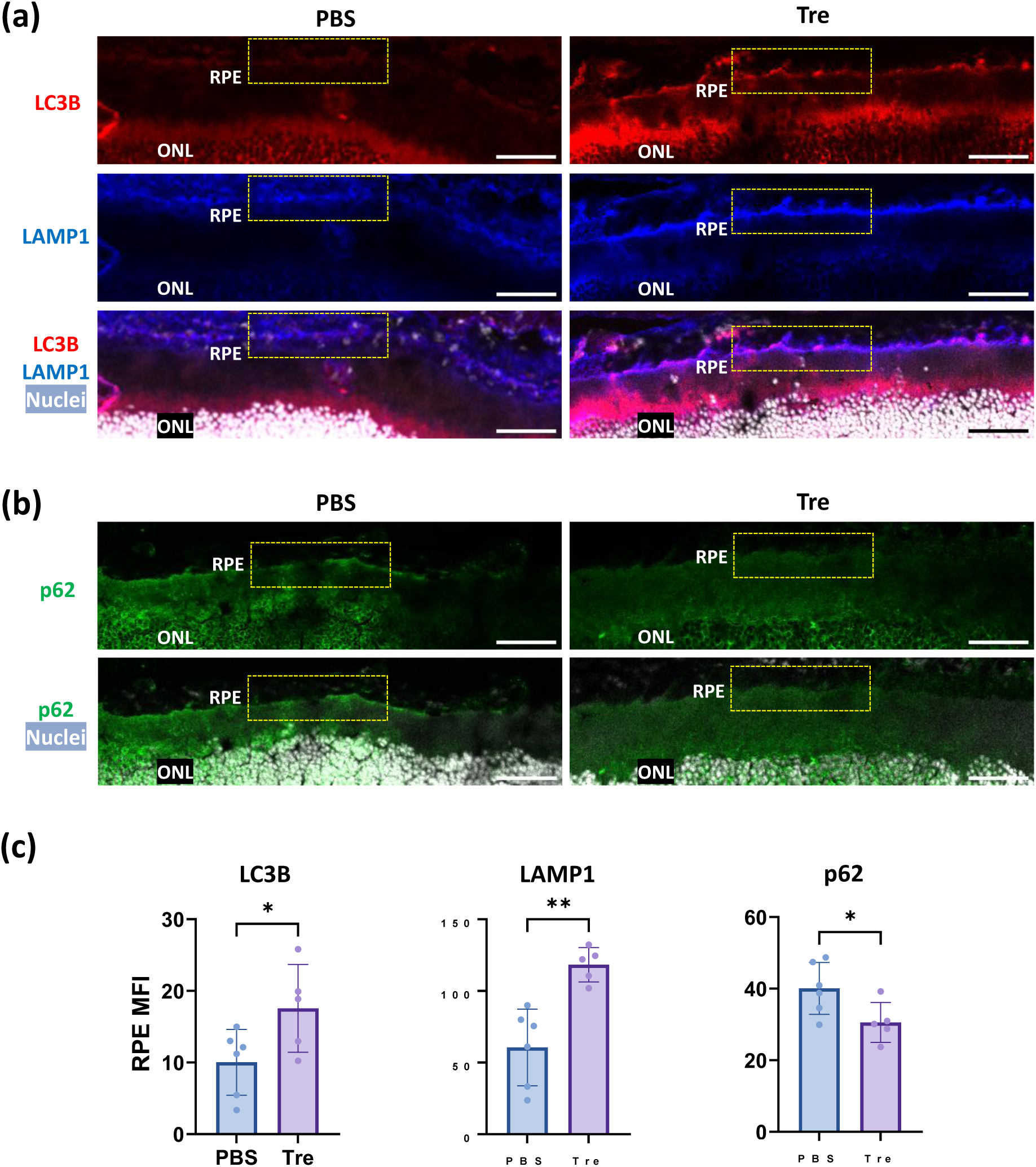
Modulation of markers of autophagy activity in murine RPE with topical trehalose application. (**a**) Representative confocal images of LC3B (red) and LAMP1 (blue) staining in retinal sections. Merge channel shows nuclei in grey, and colocalization between LC3B and LAMP1 in magenta at the RPE layer. Boxed areas indicate part of the RPE region of interest. (**b**) Representative confocal images of p62 (green) staining in retinal sections. Merge channel shows nuclei in grey. Scale bar = 50 µm. (**c**) Quantification for the mean fluorescence intensity (MFI) of LC3, LAMP1 and p62 signals in 3 different fields at the RPE region from each section (PBS: n = 6; Tre: n = 5). Comparison by Student’s t-tests for LC3B, LAMP1 and p62 (all normal data with equal variances) (**c**). *P < 0.05; **P < 0.01.

### 2.7 Topical trehalose suppresses RPE and photoreceptor loss in a light-induced retinal degeneration model

Having confirmed that topical trehalose enhances autophagic activity in murine RPE, we proceeded to evaluate its protective potential in a subacute stress model of light-induced retinal degeneration (LIRD). This model, characterized by oxidative stress and damage to the outer retina (Ding, Aredo, Zhong, Zhao, & Ufret-Vincenty, 2017; J. Liu et al., 2024), has been routinely used as a suitable pre-clinical approach to interrogate pathways operative in AMD (Carozza et al., 2024).

Seven mice were pre-treated with topical trehalose (9% in 10 µl PBS), while an equal number were treated with PBS alone, administered twice daily for seven days before light exposure. The left eye of each mouse underwent fluorescein (FL)-assisted light injury in a blinded manner, with the right eye serving as the control. Seven days post-light induction, PBS-treated eyes exhibited extensive retinal damage (Figure 6a) and over 60% thinning of the outer nuclear layer (ONL, P < 0.0001, Figure 6b). In contrast, trehalose pre-treatment significantly alleviated the damage, reducing ONL loss by 53% (P < 0.001, Figure 6b). Importantly, although trehalose treatment elevated basal autophagy flux in mouse eyes (Figure 5), no discernable differences were observed in fundus or optical coherence tomography (OCT) examination, or ONL thickness in unchallenged eyes (Figure 6).

**Figure 6:**
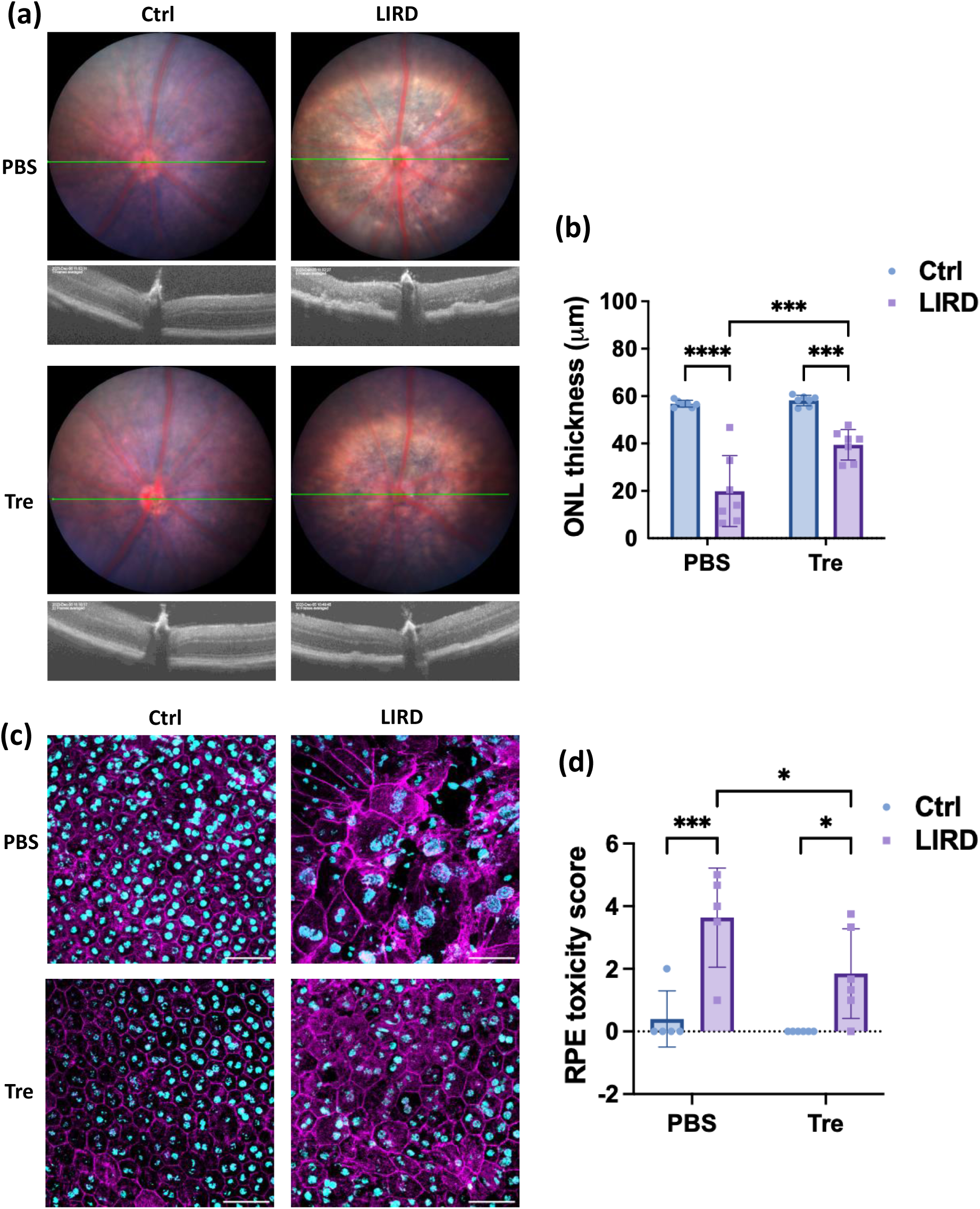
Protective effects of topical trehalose pre-treatment in a murine model of light-induced outer retinal degeneration. (**a**) Representative fundus (top) and OCT (bottom) images comparing PBS-treated (top row) and trehalose-treated (bottom row) eyes, captured 7 days post-light challenge. The green line demarks the scan line for corresponding OCT images. One-time light exposure (right column) was applied to the left eyes after 7 days of pre-treatment with either PBS or trehalose, followed by continued application for an additional 7 days before tissue harvest. Right eyes without light challenge served as undamaged controls (left column). (**b**) Quantification of averaged outer nuclear layer (ONL) thickness across the four groups (n = 7) was performed in a blinded manner. (**c**) Representative confocal images of RPE flatmounts displaying F-Actin junction staining (magenta) and nuclei (cyan). Scale bar = 50 μm. (**d**) Scoring of RPE toxicity using an established protocol in a blinded manner (n = 5). Comparisons by two-way ANOVA with Holm-Šídák tests (**b** and **d**). *P < 0.05; ***P < 0.001; ****P < 0.0001.

To assess RPE morphology and integrity, RPE flatmounts of five randomly selected mice per group in the same experimental setup were stained with Phalloidin to visualize F-actin junctions (Xiong et al., 2019). Images were captured from four representative areas 0.4 mm away from the optic nerve head (ONH) on each flatmount. As shown in Figure 6c, light-induced injury caused evident damage to the RPE, illustrated by enlarged cells, disrupted hexagonal morphology, and disarray of the F-actin structure. Topical trehalose pre-treatment preserved RPE morphology and F-actin organization (Figure 6c). Using a scoring protocol for RPE toxicity (Xiong et al., 2019), PBS-treated LIRD eyes showed a significant increase in toxicity grade, which was attenuated by trehalose pre-treatment (Figure 6d).

## 3. DISCUSSION

Here, we demonstrate a significant decline in autophagy pathways associated with RPE aging in both humans and mice, accompanied by further disruption and severe dysregulation across AMD severity stages. Furthermore, we present evidence showing the protective effects of topical trehalose in mitigating oxidative damage and light-induced degeneration in mouse RPE. This study provides important insights into the interconnected roles of aging and autophagy disruption in RPE dysfunction, supporting the potential of trehalose to enhance autophagy and restore age-related decline, thus potentially attenuate AMD progression.

Aging is a major risk factor for RPE dysfunction, driven by the tissue’s high metabolic demands throughout the lifespan (S. Wang et al., 2020). Although a direct comparison of age-associated transcriptomic changes between human RPE/choroid and mouse RPE is not fully analogous, our analyses reveal significant overlaps in downregulated pathways crucial for RPE function. These include autophagy/mitophagy, melanogenesis, stress response, mitochondrial respiration, and fatty acid metabolism (Figures 1, 3, S2, and S3). Our findings corroborate previous studies reporting age-related reductions in melanin granules, autophagic activity, and metabolic capacity in response to oxidative stress, and lipid metabolism in the RPE (Intartaglia et al., 2022; Sarna et al., 2003). Specifically, we identify a predominant downregulation of autophagy pathways with age, supporting an age-associated decline in cellular clearance mechanisms (Crabb, 2014; A. L. Wang et al., 2009). This decline leads to the accumulation of damaged organelles and macromolecule aggregates, which impair RPE function and reduce resilience to cellular stress.

In contrast, the upregulated pathways showed fewer overlaps between human and mouse samples, likely reflecting, at least in part, differences in species-specific compensatory mechanisms or stress response pathways activated during aging (Tyshkovskiy et al., 2023). For instance, the upregulation of the dilated cardiomyopathy pathway in both species supports a shared molecular link between aging RPE and cardiovascular conditions (Chang, Huang, Chou, Chang, & Sun, 2021). Furthermore, significant interspecies differences were also evident, such as the opposing trends in MAPK signaling pathway, which was downregulated in human samples but upregulated in mouse samples with age. The variations underline the need for caution when translating certain findings from mouse models to human biology for accurate modeling.

While AMD is closely linked to aging, our study uncovers previously unrecognized distinctions in autophagy-lysosome transcriptomic changes in the RPE/choroid between aging and AMD. Although aging is predominately associated with a downregulation in autophagy and lysosome pathways, AMD exhibits stage-specific regulation (Figure S4). In eAMD, there were nonsignificant changes in autophagy-related transcripts and significant bidirectional regulation in lysosome-related transcripts, suggesting a partial but insufficient compensatory response that may be undermined by age-related decline. As eAMD progressing to iAMD, the disease was characterized by pronounced upregulation of both autophagy and lysosome pathways, which we predict as an enhanced compensatory response by surviving cells to increased cellular stress and waste accumulation. However, in advanced AMD stages (GA and nAMD), the dysregulation of autophagy and lysosome pathways became more pronounced and bidirectional in general, likely due to a loss of control mechanisms and excessive decline in late-stage degenerative cells, further exacerbating tissue damage. The data highlights the critical importance of considering timing and dosing for treatments targeting autophagy in relation to AMD progression to maximize benefits and minimize potential side effects (Ambati & Fowler, 2012; S. Liu, Yao, Yang, & Wang, 2023). This notion is supported by evidence from Alzheimer’s disease (AD) models, where early autophagy induction conferred therapeutic benefits, whereas late treatment was less effective in offering protection in the experimental setting (Majumder, Richardson, Strong, & Oddo, 2011).

The relevance of this study is the demonstration of autophagy dysregulation as a critical factor in both aging and AMD progression, potentially contributing to the chronic processes underlying disease onset and exacerbation. Trehalose, known for its safety, efficient tissue penetration, and compatibility with non-invasive delivery, is a promising candidate for managing chronic disorders (Menzies et al., 2017). Long-term oral trehalose (2% in drinking water for 8 months) has shown significant efficacy in reducing retinal degeneration and maintaining visual function in mice with lysosomal hydrolase deficiencies (Lotfi et al., 2018). However, translating oral trehalose benefits to humans is challenging, due to enzymatic breakdown by trehalase in the gastrointestinal tract, reducing drug bioavailability and efficacy (Chen & Gibney, 2023).

While topical delivery to the eye’s posterior segment also poses challenges due to anatomical barriers, preclinical studies suggest retinal delivery of small molecules is achievable (Rodrigues et al., 2018). Drug penetration to posterior tissues occurs via two major routes: the corneal and conjunctival/scleral routes. However, much of the applied drug is eliminated into the systemic circulation (Rodrigues et al., 2018), with less than 3% reaching the aqueous humor and even smaller amounts accessing the posterior segment. For hydrophilic drugs including trehalose, lateral diffusion through conjunctival and sclera, followed by penetration through Bruch’s membrane and RPE, is a plausible pathway for posterior delivery (Hughes, Olejnik, Chang-Lin, & Wilson, 2005). These insights highlight the need of optimizing topical therapies for retinal diseases.

Several limitations of this study exist. First, caution is needed when making interspecies comparisons. RPE cells from young (5 months) and aged (22 months) mice were analyzed, equivalent to human ages of 20-30 and 70-80 years, respectively, but human samples (aged 50-94 years) lacked young individuals, potentially leaving certain aspects of the comparison inaccurate. Second, mouse tissues consisted of pure RPE cells, while human data included RPE/choroid, introducing a confounding factor as choroidal endothelium is also susceptible to autophagy-related oxidative damage. This difference may contribute to interspecies transcriptomic variations. Third, while trehalose’s therapeutic effects were demonstrated in mice, the extent to which enhanced autophagy protected against oxidative insults remains unclear. Constrained by the scope of the study, the impact of trehalose on other critical retinal pathology mechanisms, such as inflammation, was not investigated.

## 4. CONCLUSION

This study reveals that the age-associated transcriptomic downregulation of autophagy pathways may predispose the RPE to a loss of homeostasis and function. The relevance lies in the link of such dysregulation to AMD progression, potentially contributing to disease onset and development. Trehalose demonstrates significant potential to enhance autophagy flux and confer protection against subacute stress in a light-induced retinal degeneration model. Our findings support the therapeutic potential of targeting autophagy to protect against RPE damage and attenuate AMD progression. Future studies on trehalose’s protective effects for AMD should prioritize: 1) the use of aging and chronic models to more align with human disease pathology; 2) larger animal models to validate translational potential; and 3) sustained-release formulations or advanced drug delivery techniques to enhance drug retention, durability, and penetration to the RPE and outer retina.

## 5. MATERIALS AND METHODS

### 5.1 Human transcriptomic data mining

Human RNA-seq datasets (Orozco et al., 2023) were used for RPE/choroidal analysis. The study included mixed-gender subgroups of 36 normal controls (mean age 70.5 ± 13.6 years), 16 eAMD (80.1 ± 10.0), 8 iAMD (85.1 ± 6.4), 10 GA (86.8 ± 5.8), and 18 nAMD (80.7 ± 11.4). DEGs significantly correlated with increasing age, or with dry (dAMD) progression from eAMD to GA identified via linear analysis, and stage-specific DEGs for AMD compared to normal controls, were extracted from the original study.

Pathway enrichment of DEGs was performed using Metascape (Zhou et al., 2019). The Membership Search feature was employed with specific keywords to query KEGG or Rectome pathways, to identify associations between DEGs and pathways of interest. Default parameters were used for enrichment analysis: P cutoff = 0.01, minimum overlap = 3, and minimum enrichment = 1.5.

### 5.2 Mice

C57BL/6J male mice were from Charles River Laboratories and housed in the Animal Services Unit at the University of Bristol, in accordance with the Home Office Regulations. Treatment of the animals conformed to UK legislation and the Association of Research in Vision and Ophthalmology statement. The experiments involving animals were conducted in accordance with the approved University of Bristol institutional guidelines and all experimental protocols under Home Office Project License PP9783504 were approved by the University of Bristol Ethical Review Group.

Mice for RNA-seq analysis were randomly selected and euthanized at ages of 5 and 22 months via cervical dislocation. Primary RPE cell cultures were derived from mice aged 8-10 weeks. For *in vivo* experiments assessing the effects of topical trehalose on the LIRD model, mice entered the study at 9 weeks of age and were culled after 2 weeks for tissue analysis. All tissue samples were collected 3-4 h after the onset of the light cycle (lights ON between 7:00 am and 7:00 pm) to minimize circadian influences.

### 5.3 Mouse RPE isolation

Murine RPE cells were isolated as described before (J. Liu et al., 2024; Scott et al., 2021). Briefly, eyes were enucleated and cleaned using angled scissors to remove any remaining connective tissue. After removing the cornea and lens, eyecups were incubated at 37°C in hyaluronidase (Sigma-Aldrich) for 45 min, and in HBSS buffer with 10 mM HEPES for a further 30 min before retinas were removed via incision. Subsequently, eyecups were incubated at 37°C in trypsin/EDTA for 45 min before being transferred into HBSS with 20% heat-inactivated FCS. Gently shaking facilitated the detachment of RPE sheets, which were incubated in trypsin/EDTA for 1 min to form single-cell suspensions. The cells were either processed for RNA extraction or resuspended in cell culture medium for cultivation and subsequent treatments as specified. RPE purity was confirmed via immunoblotting for RPE65 and rhodopsin as demonstrated before (Scott et al., 2021), and supported by PaGenBase analysis for cell specificity of DEGs.

### 5.4 RNA-seq and analysis

RNA of RPE cells was extracted using RNeasy Mini kit (Qiagen). RNA samples were processed in the Genomics Facility at University of Bristol for ScreenTape assessment to ensure RNA integrity (RIN) > 7. Library preparation was conducted using Illumina Stranded Total RNA Prep with Ribo Zero Plus (Illumina), followed by sequencing using the Illumina NextSeq 550 platform with 2×75 bp by paired-end reads at a depth of 16 million reads.

Galaxy software was used to assess the quality of reads via FastQC, followed by read trimming via Trimmomatic. Reads were then aligned using HISAT2 and quantified using FeatureCounts. Batch correction of the resulting counts from two independent experiments was performed using R Studio v2023.06.1 with ComBat-seq. Following correction, counts were normalized by DESeq2 package and lowly expressed genes (basemean < 20) were removed. Normalized data of the remaining 14220 genes were processed using iDEP.96 for logarithmic transformation, quality control, and DEG identification (Ge, Son, & Yao, 2018). P values were adjusted using the Benjamini–Hochberg method. DEGs were selected based on an adjusted P value (Padj) < 0.05 and |log2FC| > 0.25.

Cell and tissue specificity analysis, along with pathway enrichment of DEGs, were conducted using Metascape platform with the PaGenBase and KEGG databases, respectively. Membership analyses were applied with specified KEGG or Rectome pathway terms.

### 5.5 Primary murine RPE cell culture

Isolated murine RPE cells were resuspended in alpha MEM supplemented with 1% N1 Medium Supplement (Sigma-Aldrich), 1% L-glutamine, 1% penicillin–streptomycin, 1% nonessential amino acid solution (Thermo Fisher Scientific), 20 μg/l hydrocortisone (Sigma-Aldrich), 250 mg/l taurine (Sigma-Aldrich), 0.013 μg/l triiodo-thyronin (Sigma-Aldrich), and 5% FCS. The cells were seeded to laminin (Sigma-Aldrich)-precoated 24-well cell culture plates, Seahorse XFp Cell Culture Plates (Agilent Technologies), or 16-well chambered cover glass (Thermo Fisher Scientific), at a density of 25,000/cm^2^. Serum was withdrawn after one week of incubation, and the cells were kept in serum-free condition for an additional week for specified treatments. Cell culture media were refreshed twice weekly.

### 5.6 Cytotoxicity assay

RPE cells were incubated with different concentrations of trehalose (6.25, 12.5, and 25 mM, Sigma-Aldrich, PHR1344) for 24 h. Cell culture supernatants were collected for assessment using a Lactate Dehydrogenase (LDH) cytotoxicity kit (Abcam).

### 5.7 Seahorse metabolic assay

Effects of trehalose and/or H_2_O_2_ on mitochondrial respiratory in RPE cells were assessed using Mito Stress tests on a Seahorse Cell Metabolic Analyzer (Agilent Technologies). Seahorse XFp cell culture miniplates, sensor cartridges, and all reagents were from Agilent Technologies. Cells were incubated in Seahorse XF DMEM containing 25 mM glucose, 1 mM pyruvate, and 2 mM glutamine in 37°C incubator without CO_2_ for 45 min. Oligomycin (1 μM), Carbonyl cyanide-p-trifluoromethoxyphenylhydrazone (FCCP, 0.5 μM) and antimycin A/rotenone (1 μM) were injected where indicated. OCR (pmol O_2_/min) was measured in real-time and normalized by total protein analyzed using a BCA assay. OCR parameters were calculated using the following formulae: nonmitochondrial respiration (NMR, minimum OCR after antimycin A/rotenone injection), basal respiration (BR, difference between OCR before oligomycin and NMR), maximal respiration (MR, difference between maximum OCR after FCCP injection and NMR), spare respiratory (SR, difference between MR and BR), ATP production (difference between OCR before oligomycin injection and minimum OCR after Oligomycin).

### 5.8 Autophagy flux measurement

The formation of autophagosome and autolysosome in RPE cells was monitored through LC3B localization using a Premo™ Autophagy Tandem sensor Kit (Thermo Fisher Scientific), which detects LC3B positive, neutral pH autophagosomes in green fluorescence (GFP) and LC3B positive, acidic pH autolysosome in red fluorescence (RFP). The reagent was added to RPE cells (40 particles/cell). After 24 h, cells were treated with trehalose (12.5 mM) for an additional 16 h. Subsequently, 50 nM of LysoTracker Deep Red (Thermo Fisher Scientific) was applied to the cells for 30 min for lysosome staining. After washing with PBS and counter-staining with Hoescht 33342 (Thermo Fisher Scientific), cells were imaged live on a Leica SP5II confocal laser scanning microscope. Z-stack images of cells were acquired using 1-µm step size and visualized using maximal intensity projections. LC3B-positive vacuoles were quantified using Fiji.

### 5.9 Mitochondrial superoxide staining

To detect mitochondrial superoxide in primary murine RPE cells treated with trehalose and/or H_2_O_2_, MitoSOX Red (Thermo Fisher Scientific) was added to the cells at a final concentration of 5 μM for 10 min. After washing with HBSS, cells were stained with Hoechst 33342 and observed using the confocal microscope.

### 5.10 Topical administration of trehalose

Trehalose (Sigma-Aldrich, T9449) powder was dissolved in sterile PBS to create final concentrations of 3% or 9% w/v (87.6 or 262.8 mM) for topical application. For safety assessment, 10 µl of the trehalose solution was applied to one eye, with an equal volume of PBS applied to the contralateral eye. In the light-induced retinal degeneration (LIRD) model for functional testing, 10 µl of high-dose trehalose or PBS was applied to both eyes twice daily at a 6-h interval, for 7 days prior to the light challenge. Trehalose treatment was paused on the day of light challenge, and resumed for an additional 7 days, after which the mice underwent clinical assessment and were sacrificed for tissue analysis.

### 5.11 Light-induced retinal degeneration

Outer retinal degeneration in mice was induced by fluorescein (FL)-assisted light-induced oxidative damage as described before (J. Liu et al., 2024). Briefly, mice were dark-adapted overnight and administered intraperitoneally with 100 µL of 2% FL (Huddersfield Pharmacy Specials) in sterile stilled water. 3 min after the injection, light was delivered at the centre of the left eye at an intensity of 18 kLux for a one-time exposure of 5 min, guided by the fundus camera of a Micron IV device (Phoenix Research Labs). The right eye was left as a control. The experimenter performing the light challenge was blinded to the treatment conditions. Following light challenge, mice were kept under normal lighting conditions.

### 5.12 Fundoscopy and optical coherence tomography

Pupils were dilated using topical eyedrops containing 1% tropicamide and 2.5% phenylephrine, and the mice were anaesthetized using 2% isoflurane inhalation. The Micron IV retinal imaging microscope (Phoenix Research Laboratories) was used to capture OCT scans and brightfield fundal images, with a gain of 3 dB and an FPS of 15. OCT scans were taken in 30-degree increments, centered at the optic disc. The ONL thickness was measured on each OCT image across a 400-µm span, beginning 100 µm from either side of the optic nerve head (ONH), resulting in a total of 16 measurements per eye to calculate an average. An independent investigator conducted the measurements in a blinded manner.

### 5.13 Immunohistochemistry and fluorescence staining

To examine the expression of autophagy and lysosome-related proteins in mouse retinal sections, eyes were enucleated and serial cryosections of 12-µm thickness were prepared using a cryostat. Sections were fixed with 4% paraformaldehyde (PFA) for 15 min and washed thrice before blocking with 5% bovine serum albumin (BSA), 5% normal donkey serum (NDS) and 0.2% Triton X-100 in PBS. Sections were stained overnight at 4°C with primary antibodies diluted in staining buffer (1% BSA in PBS), containing rabbit anti-LC3B (1:1000, Abcam, ab5152), rat anti-LAMP1 (1:50, Santa Cruz Biotechnology, sc-19992), or rabbit anti-p62 (1:1000, Sigma-Aldrich, P0067). After washing, samples were incubated with donkey anti-rabbit IgG-Alexa Fluor (AF)488 or anti-rat IgG-AF555 (1:400, Thermo Fisher Scientific) for 1 h. Slides were then counter-stained with Hoescht 33342. After mounting with anti-fade fluorescence mounting medium (Abcam), samples were imaged with a Leica SP5II confocal laser scanning microscope focused on the RPE region of interest. Mean fluorescence intensity (MFI) was quantified using Fiji.

To prepare RPE/choroid wholemounts for assessment of RPE morphology, enucleated eyes were dissected and RPE-choroid-sclera isolated. Tissues were fixed in 4% PFA for 1 h, washed in PBS, and blocked and permeabilized with 5% BSA, 5% NDS, and 0.3% Tween-20 for 2 h. Tissues were then incubated with Phalloidin-AF555 (1:20 in PBS, New England Biolabs) for 30 min to stain F-actin. After washing and staining with Hoescht 33342, samples were mounted and imaged using the confocal microscope.

### 5.14 Statistics

Statistical methods for RNA-seq analysis were described elsewhere. All other statistical analyses were conducted using GraphPad Prism 9.0. For data analysis on experiments with one variable, normality of samples was determined by Shapiro-Wilk test, and homoscedasticity across groups with normal data was measured via F-test. Comparison between two groups with a single variable was performed using unpaired two-tailed Student’s t-test for normal data with equal variances (Figure 5c), Welch’s test for normal data with unequal variances (Figure 4b), or Mann-Whitney test if at least one group has nonnormal data (Figure 4b). For experiments involving more than two groups and one variable, comparisons were measured using one-way ANOVA followed by Bonferroni post-hoc multiple comparisons if all groups are normal and equal (Figure S5a). In experiments involving two independent variables, we used two-way ANOVA with Bonferroni tests for *in vitro* experiments (Figure 4d, and Figure S5c), while with Holm-Sidak tests for *in vivo* experiments to enhance statistical power and minimize the need for additional animals (Figure 6b,d). Differences between groups were considered significant at P < 0.05. Results are presented as means ± standard deviation (SD).

## ACKNOWLEDGMENTS

The authors wish to acknowledge the assistance of the Wolfson Bioimaging Facility at the University of Bristol.

## CONFLICT OF INTEREST STATEMENT

The authors declared no conflicts of interest.

## FUNDING STATEMENT

This study was supported by the Sight Research UK (grant SAC052 to JL) and Macular Society (grant 21-RG-1 to ADD).

## AUTHOR’S CONTRIBUTIONS

K.C., N.R., K.O., J.W., and J.L. conducted the experiments and analyzed the data. G.S., Z.L, S.C., and J.L.B.P. analyzed the data. K.C., J.L., and N.R. drafted the manuscript and prepared the figures. J.L., A.D.D., L.B.N., and K.C. conceptualized the study and designed the experiments. J.L. and A.D.D. secured funding, provided supervision, and revised the manuscript. All authors reviewed, revised, and approved the final version of the manuscript.

## DATA AVAILABILITY

All data are available in the main text or the supplementary materials. The murine datasets used for this study can be found in the Gene Expression Omnibus (GEO) repository under the accession number GSE284125 (access token to be provided upon request).

## SUPPORTING INFORMATION

Figures S1-S6

Data files S1-S5

## Notes

### Competing Interest Statement

The authors have declared no competing interest.

