## Supplementary figures for "Age-Associated Decline in Autophagy Pathways in Retinal Pigment Epithelium and Protective Effects of Topical Trehalose in Light-induced Outer Retinal Degeneration in Mice"

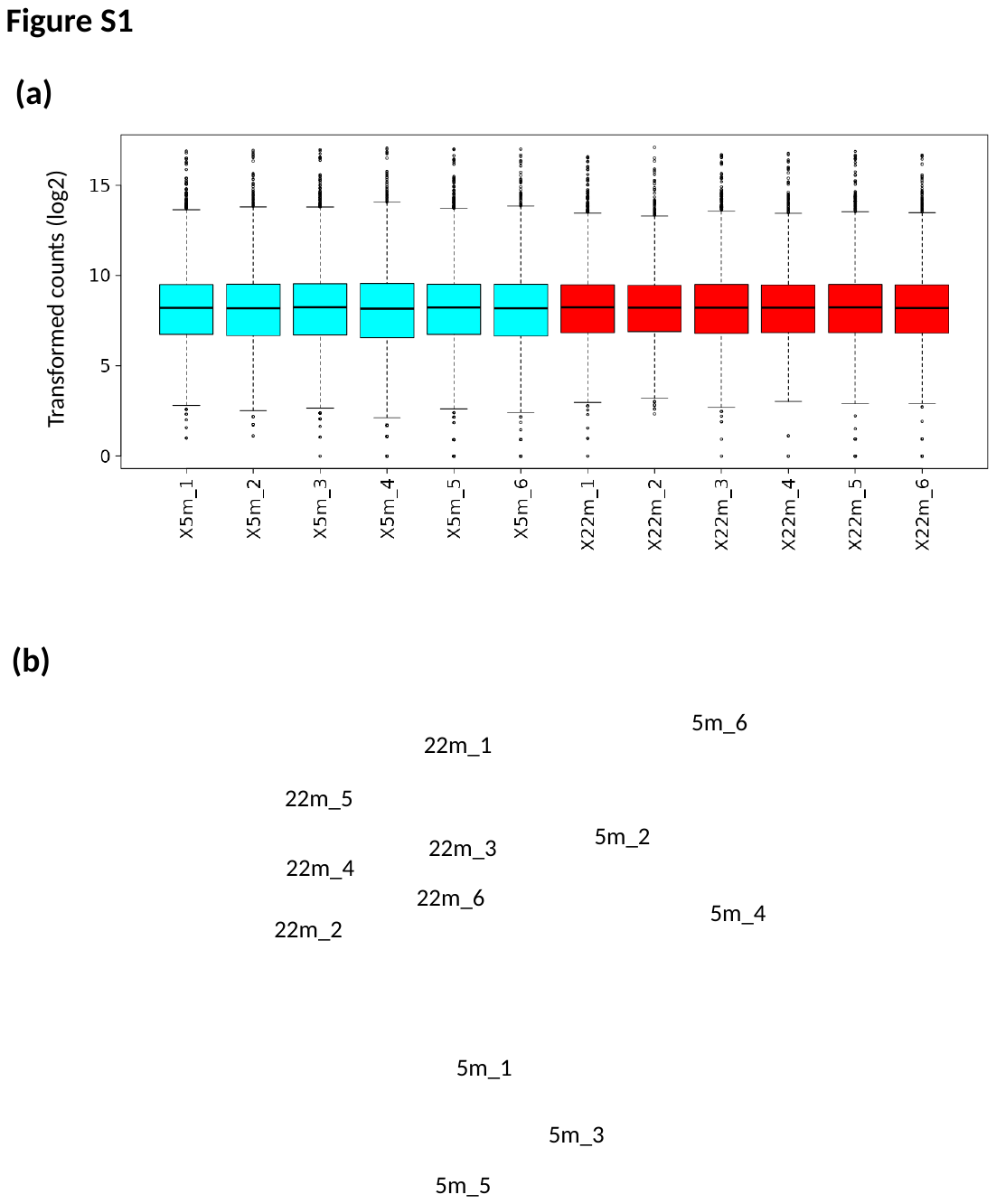

**Figure S1. Quality control of RNA-seq data from murine RPE using iDEP.96 platform.** RPE cells isolated from 5-month-old (cyan) or 22-month-old (red) male C57BL/6J mice (n = 6 per group) were subjected to RNA-seq analysis. (**a**) The box plot of normalized and transformed count showing data distribution. (**b**) The Principal Component Analysis (PCA) plot, based on expressed genes from samples, visualizing data variance. The first principal components (PC1) and the second principal component (PC2) are shown, with each data point signifying a distinct sample.

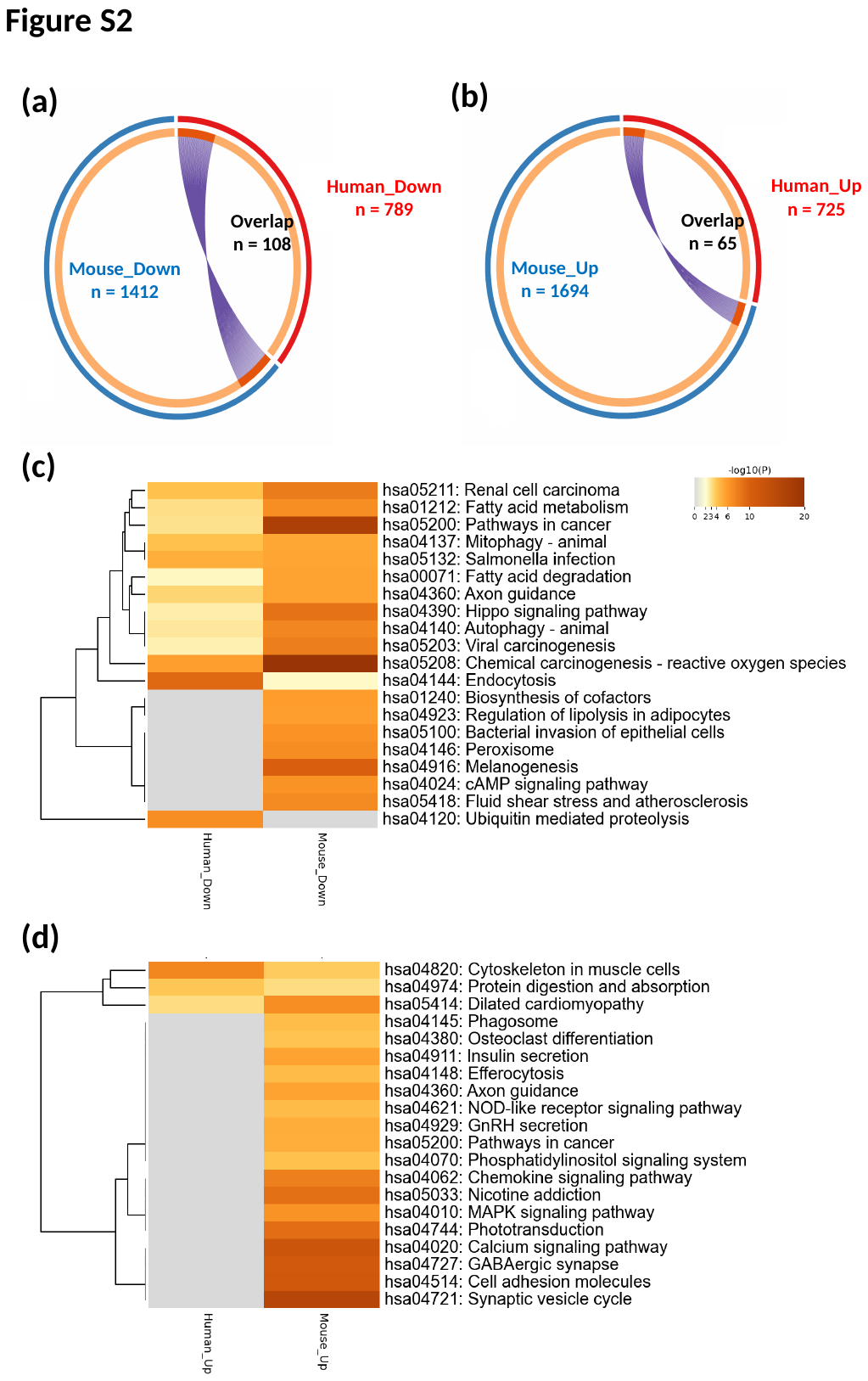

**Figure S2. Shared transcriptomic changes between aged human RPE/choroid and mouse RPE.** (**a** and **b**) Circos plots generated from multiple gene list analyses via Metascape illustrate the overlap of downregulated (**a**) and upregulated (**b**) genes between aged human (red) and mouse (blue) samples. The outer arcs represent the identity of each gene list, while the inner arcs depict the distribution of genes within each list. Dark orange indicates genes shared across multiple lists, while light orange represents genes unique to a specific list. Purple lines connect identical genes shared between lists. Numbers denote the total number of DEGs and the count of overlapping genes between human and mouse datasets. (**c** and **d**) Heatmaps of enriched KEGG pathways associated with downregulated (**c**) and upregulated (**d**) genes in human and mouse datasets.

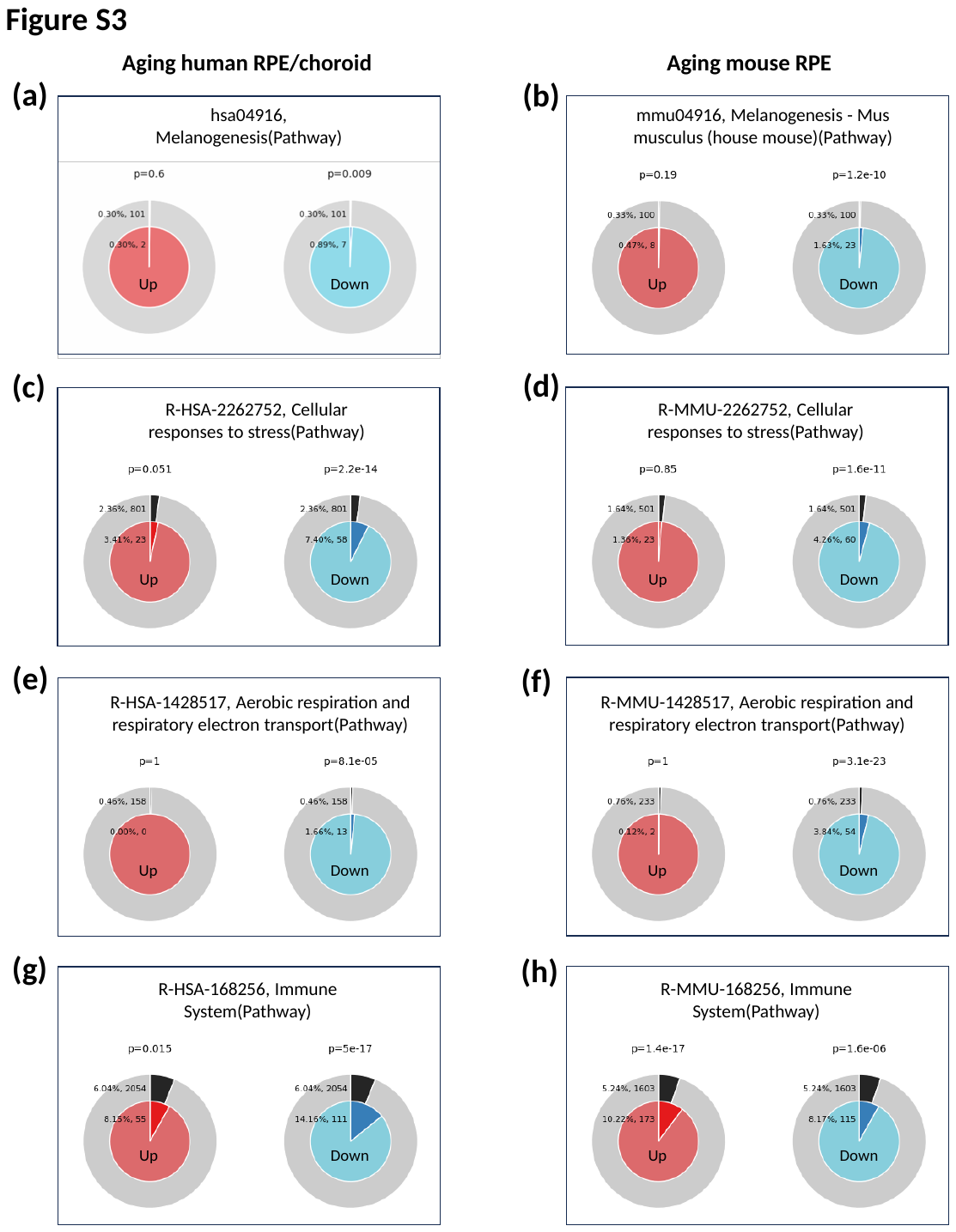

**Figure S3. Comparative transcriptomic changes between aged human RPE/choroid and mouse RPE using Membership analysis.** DEGs from human (**a**, **c**, **e**, and **g**) and mouse (**b**, **d**, **f**, and **h**) samples were analyzed using Membership analysis via the Metascape platform. Enrichment of upregulated (left) and downregulated (right) genes associated with specific KEGG or Reactome pathway terms, including Melanogenesis (human: hsa04916, **a**; mouse: mmu04916, **b**), Cellular response to stress (human: R-HSA-2262752, **c**; mouse: R-MMU-2262752, **d**), Aerobic respiration and respiratory electron transport (human: R-HSA-1428517, **e**; mouse: R-MMU-1428517, **f**), and Immune System (human: R-HSA-168256, **g**; mouse: R-MMU-168256, **h**). The outer circle represents the percentage and number of total genes associated with the pathway, while the inner circle shows the percentage and number of input genes mapping to the pathway. The P value above the pie chart indicates the statistical significance of the gene set membership to the pathway term.

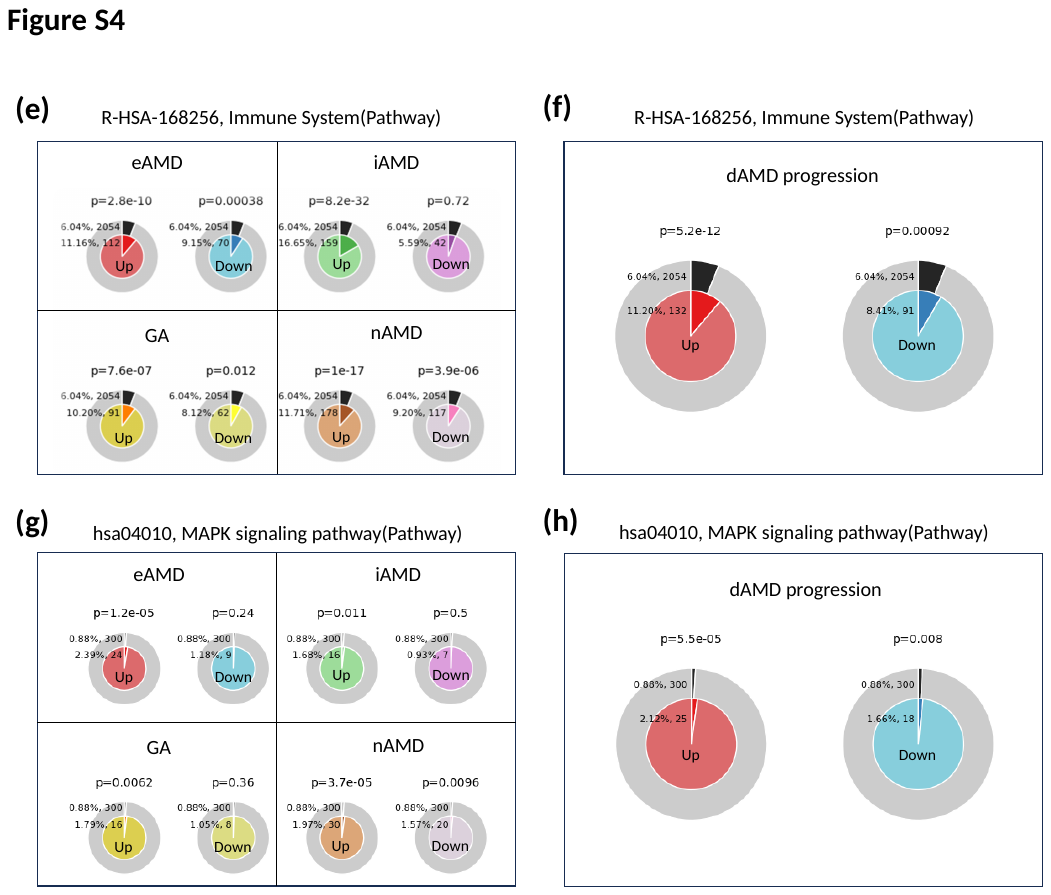

**Figure S4. Dynamic alterations in selected transcriptomic pathways across AMD stages.** (**a**, **c**, **e, and g**) Metascape-based Membership analysis of a bulk RNA-seq dataset (Orozco et al. 2023) on specific transcriptomic pathways across individual stages of dry AMD (dAMD) and neovascular AMD (nAMD). eAMD: early AMD; iAMD: intermediate AMD; GA: geographic atrophy. (**b**, **d**, **f**, and **h**) Membership analysis of the same dataset on corresponding pathways during the progression of dAMD based on the linear analysis of dAMD stages. Enrichment of upregulated (left) and downregulated (right) genes associated with specific KEGG or Reactome pathway terms, including Autophagy (hsa04140, **a** and **b**), Lysosome (hsa04142, **c** and **d**), Immune System (R-HAS-168256, **e** and **f**), and MAPK signaling pathway (hsa04010, **g** and **h**). The outer circle indicates the percentage and number of total genes associated with the pathway term, and the inner circle represents the percentage and number of input genes mapping to the term. The P-values above the pie charts denote the statistical significance of the association between the input gene set and the pathway term.

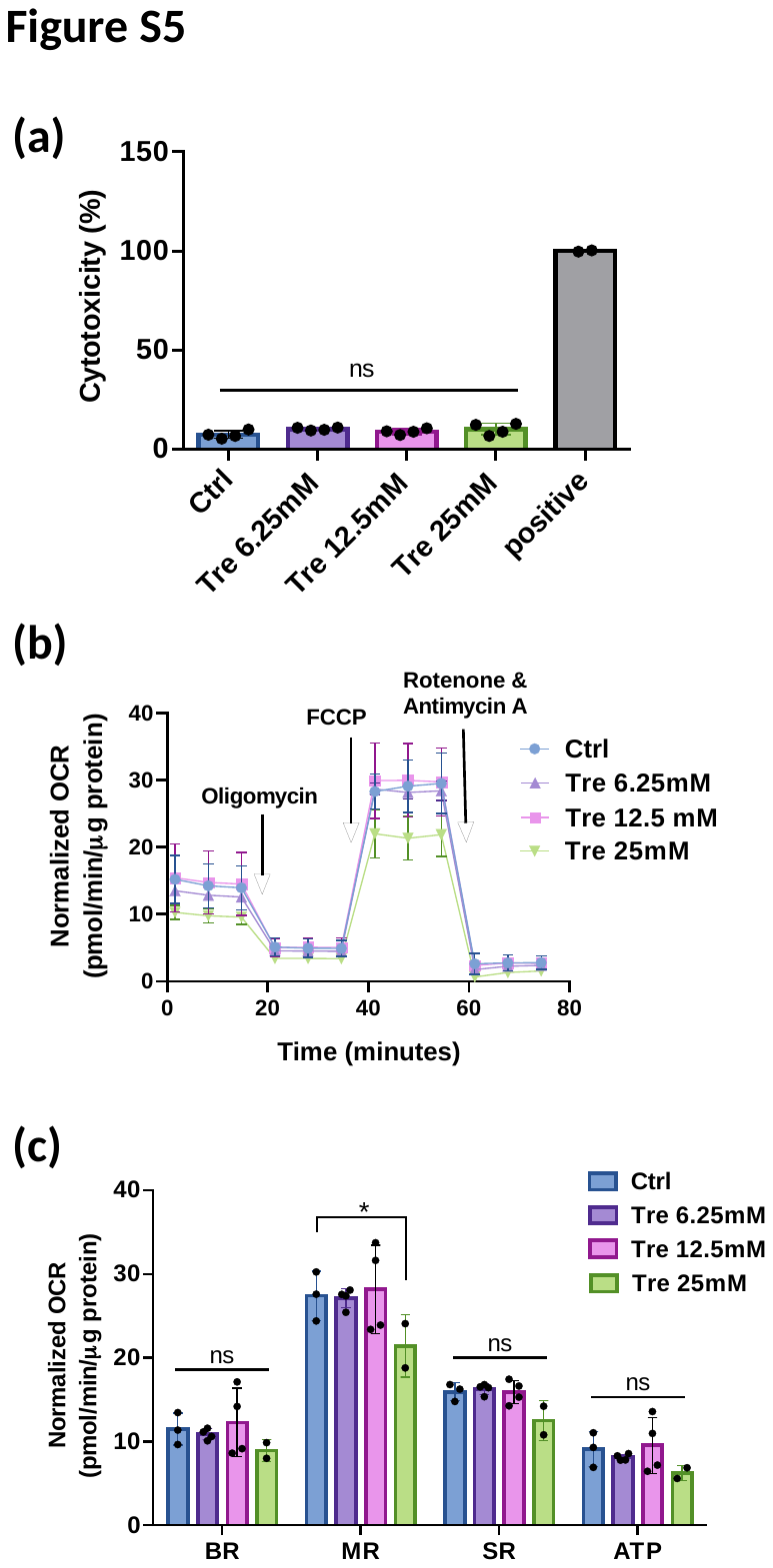

**Figure S5. Identification of safe dose of trehalose in primary murine RPE cells *in vitro*.** (**a**) LDH cytotoxicity analysis of primary RPE cells post 24-hour treatment with different concentrations of trehalose. The RPE cell lysate was used as a positive control (100% toxicity). (**b** and **c**) Mito stress analysis of primary RPE cells treated with the same conditions as (**a**). OCR profile (**b**) and parameters (**c**) of the cells were shown. BR: basal respiration; MR: maximal respiration; SR: spare respiration; ATP: ATP production. *P < 0.05; ns, nonsignificant. Comparison by one-way ANOVA (**a**, n = 2-4) or two-way ANOVA (**c**, n = 2-4), both with Bonferroni post-hoc tests.

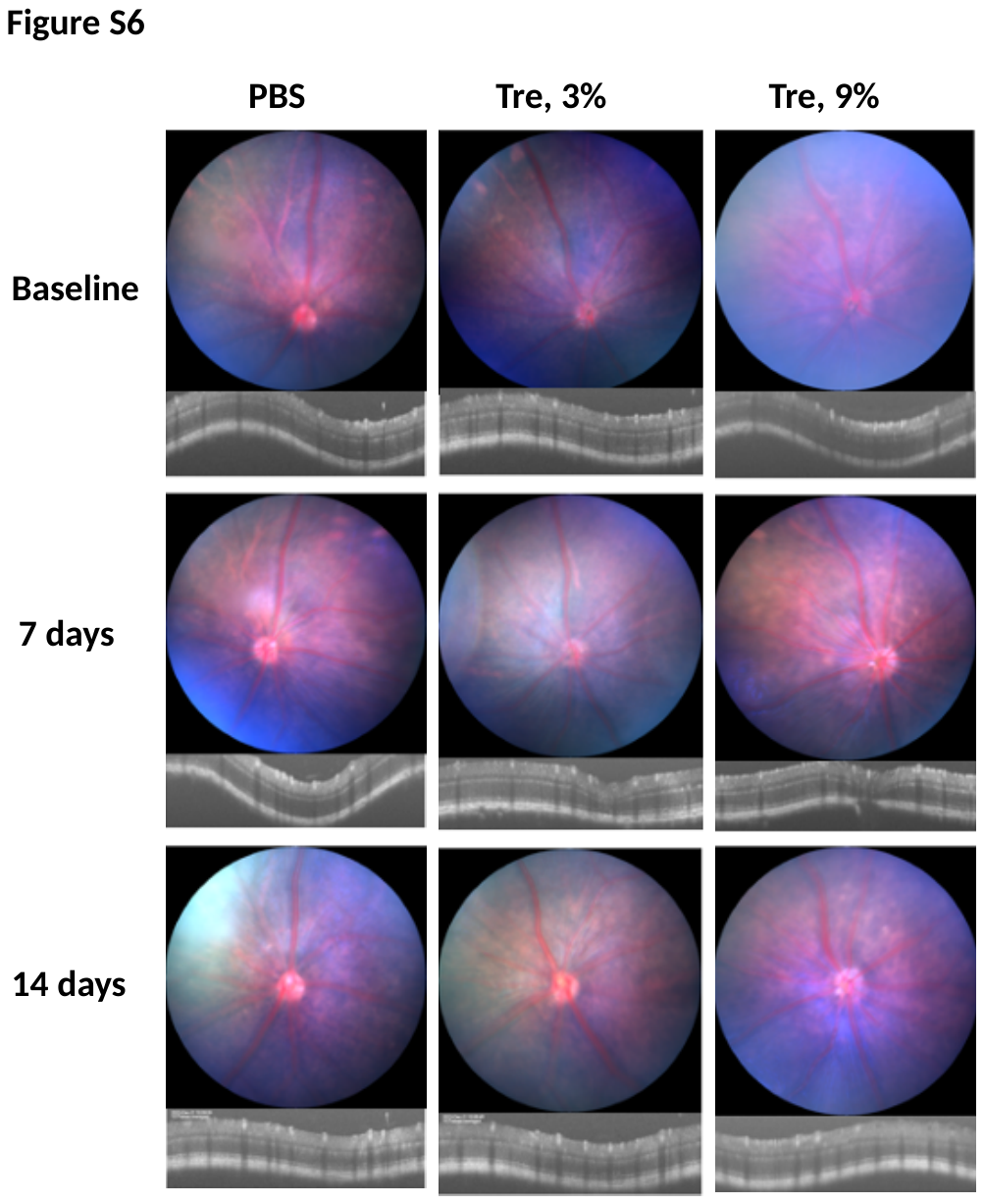

**Figure S6. No significant side effects with topical trehalose application.** Representative fundal and OCT images of mice were captured at baseline (before treatment) and at 7 and 14 days following twice-daily administration (with a 6-hour interval) of 10 µl of either PBS (left) or trehalose solutions (3%: middle; 9%: right) dissolved in PBS. Images are representative of 4 eyes per group.
